# Recognizing off-sample mass spectrometry images with machine and deep learning

**DOI:** 10.1101/518977

**Authors:** Katja Ovchinnikova, Vitaly Kovalev, Lachlan Stuart, Theodore Alexandrov

## Abstract

**Motivation:** Imaging mass spectrometry (imaging MS) is a powerful technology for revealing localizations of hundreds of molecules in tissue sections. However, imaging MS data is polluted with off-sample ions caused by caused by sample preparation, particularly by the MALDI matrix application. The presence of the off-sample ion images confounds and hinders metabolite identification and downstream analysis.

**Results:** We created a high-quality gold standard of 23238 manually tagged ion images from 87 public datasets from the METASPACE knowledge base. We developed several machine and deep learning methods for recognizing off-sample ion images. Deep residual learning performed the best with the F1 score of 0.97. Spatio-molecular biclustering method achieved the F1 scores of 0.96 and 0.93 in semi- and fully-automated scenarios, respectively. Molecular co-localization method achieved the F1 score of 0.90. We investigated the clusters of the DHB matrix, the most common MALDI matrix, and characterized parameters of a clusters combinatorial model. This work addresses an important issue in imaging MS and illustrates how public data, modern web technologies, and machine and deep learning open novel avenues in imaging MS.

**Availability and Implementation:** Data and source code are available at: https://github.com/metaspace2020/offsample.

## Introduction

Imaging mass spectrometry (imaging MS) emerged as a powerful and versatile technology for spatial molecular analysis (Doerr, 2018; Dreisewerd and Yew, 2017; Buchberger *et al.*, 2018) with a particular interest in clinical and pharma applications (Vaysse *et al.*, 2017; Schulz *et al.*, 2018). The capacities and potential of imaging MS was boosted with the introduction of high- and ultrahigh-resolving power mass spectrometry analyzers (FTICR and Orbitrap) that rapidly shifted the current focus towards applications in metabolomics and lipidomics (Palmer *et al.*, 2016). Among various flavors of imaging MS, matrix-assisted laser-desorption ionization MALDI-imaging is the most widespread (Palmer *et al.*, 2016). MALDI-imaging requires applying the so-called matrix onto the tissue section leading to formation of a layer of crystals covering the sample to facilitate “soft desorption” energy transfer to sample analytes, enhance their ionization, and reduce in-source fragmentation (Karas and Krüger, 2003). However, the addition of ionizable matrix molecules contaminates the data with matrix ion signals which are not relevant for the molecular content of the sample and, moreover, can be isomeric or isobaric with sample analytes and thus mask out their spatial distribution. The matrix signals cannot be eliminated from the data with a simple approach, e.g. by considering [M+H]+ matrix ions since a matrix forms hundreds of so-called matrix clusters, namely ions composed of matrix molecules. The presence of matrix cluster ions pollutes MALDI-imaging data by adding biologically irrelevant signals. This potentially leads to false hits in metabolite identification and introduces confounding effects for the downstream analysis that hinders data classification, quality control, spatio-temporal studies, or pathways analysis. Despite these negative effects being widely recognized, the formation of the matrix clusters in general, as well as for specific MALDI matrices, sample preparation protocols, and types of samples, is poorly understood. A combinatorial model for matrix clusters was proposed (Keller and Li, 2000) which, however, suggests thousands of theoretically possible clusters whereas only some of them are observed in real experiments.

Matrix ion images can be recognized after visual expert examination. A typical matrix image normally exhibits the so-called “background” or “off-sample” pattern with high intensities in the area outside of the sample and low intensities within the sample area due to the ion suppression effect (Lee *et al.*, 2014; Taylor *et al.*, 2018). However, manual filtering of off-sample ion images is often not feasible due to the sheer amount of ion images: 10^4^−10^7^ in an imaging MS dataset, 10^2^−10^3^ after a molecular annotation (Palmer *et al.*, 2017). To the best of our knowledge, no automated or semi-automated methods for recognizing off-sample ion images were published. The software MSiReader (Bokhart *et al.*, 2018) provides a semi-automatic method that requires manual selection of the off-sample area and a threshold for finding ion images co-localized within this area; no evaluation or error analysis of this method was published however. Other software packages also provide co-localization analysis methods which can be used for pattern matching but were never investigated specifically for off-sample image recognition. We explain the lack of methods for off-sample image recognition as being due to: a high spatial heterogeneity of the signals both for the sample and matrix itself, diversity of shapes of samples, low signal-to-noise ratio, and high computational requirements due to a high number of ion images. Moreover, the variety of samples, matrix-application protocols, MALDI matrices used in MALDI-imaging requires a robust method applicable in a wide range of scenarios. Note that other types of imaging MS (e.g. DESI-imaging) are also prone to the issue of generating background ion images; see e.g. (Calligaris *et al.*, 2014). We believe that the key factor preventing computational developments to address this gap is the lack of annotated data for evaluation of potential recognition methods.

We recently developed a bioinformatics method for metabolite and lipids annotation in imaging MS (Palmer *et al.*, 2017). This method reduces millions of m/z-channels in imaging MS data to hundreds to thousands of images associated with molecules from a molecular database of interest. We implemented the method as a cloud software platform called METASPACE (http://metaspace2020.eu) and have received more than 3000 submissions of imaging MS data from over 50 labs, representing various biological systems analyzed using various sample preparation techniques and MALDI matrices. The images from over 2800 public submissions represent an open community-populated knowledge base of metabolic ion images.

In this paper, we tackled the issue of recognizing off-sample ion images by capitalizing on the information contained in the public METASPACE knowledge base. We used public METASPACE datasets to study the problem and to create a gold standard set of off-sample ion images tagged by a team of experts. We developed several algorithms for recognition of off-sample images. We evaluated them on the gold standard data and performed error analysis. We applied the developed methods to investigate which matrix clusters are most commonly observed in MALDI-imaging when using one of the most popular matrices, 2,5-dihydroxybenzoic acid (DHB) (Strupat *et al.*, 1991).

## Methods

### Creating a gold standard set of ion images

In image recognition, a gold standard set is a collection of images manually tagged by experts called taggers often curated to ensure the highest quality. Having a gold standard set enables training and evaluating machine learning algorithms. However, creating an unbiased, representative, and balanced gold standard is a substantial challenge on its own. To the best of our knowledge, there exists no gold standard set of off-sample images for imaging MS.

#### Selecting imaging MS datasets for the gold standard

To create a gold standard set of off-sample and on-sample ion images, we selected public datasets from METASPACE (http://metaspace2020.eu) with the aim to have a diverse yet representative sample of datasets collected by different labs, for different samples, and with different imaging MS methods. At the same time, we considered a manageable number of datasets for manual tagging of ion images as every dataset in METASPACE contains 10^1^-10^3^ ion images.

First, we randomly selected 100 datasets from the public datasets in METASPACE. We added 14 more manually picked datasets to increase the diversity of the set and to represent a particular lab, biological model, sample preparation protocol, imaging MS source or analyzer that were not picked by the random sampling. We excluded 27 datasets from one big METASPACE submitter to reduce the lab, technology, and sample type bias, which left us with 87 gold standard imaging MS datasets.

#### Pilot study

Before creating a gold standard, we ran a pilot study to investigate the difficulty of recognizing off-sample ion images, as well as to learn potential pitfalls and obstacles of the tagging process. In the pilot study, we involved two taggers, PilotTagger1 and PilotTagger2. Each tagger used their own semi-automated strategy to tag ion images.

PilotTagger1 selected 19 public datasets from METASPACE. For each dataset, PilotTagger1 semi-automatically outlined the off-sample area by using a k-means clustering-based spatial segmentation. For each ion image, the tagger scaled its intensities to [0,1] and calculated the sum of the intensities within the off-sample area. Then, the tagger selected ion images with the highest values of this score as off-sample, and ion images with the lowest values of this score as on-sample.

PilotTagger2 selected 110 public datasets from METASPACE. For each dataset, PilotTagger2 picked a template ion image for the off-sample area and, for all other ion images, calculated the Spearman correlation with the template ion image. Images with the correlation above a manually selected threshold were selected as off-sample; images with the similarity score below a manually selected threshold were selected as on-sample. Based on their expert knowledge, PilotTagger2 curated all selected images by opening them in a file browser and removing those which looked to be wrongly assigned.

#### Web app for manual tagging of ion images

After the pilot study, we made the decision that to create a high-quality gold standard set of ion images, every ion image should be tagged manually, since pre-selection by an algorithm introduces a bias. We also realized that tagging requires a special tool that would not only facilitate tagging but would also allow one to inspect tagged images, correct the tagging, and curate the gold standard by another expert. Taking into account the complexity of these tasks and the lack of existing solutions, we developed our own web app TagOff for tagging ion images either as off-sample or on-sample (available at https://github.com/metaspace2020/offsample). For a public dataset in METASPACE, the TagOff web app downloads ion images from METASPACE using the GraphQL API (<u>link</u>), shows ion images, and let a tagger tag each image into one of three classes (“off-sample”, “on-sample”, “unknown”). To facilitate the tagging, we implemented several tagging either by a single- or double-mouse click or, alternatively, by pressing shortcut keys while hovering the mouse. Also, we implemented a random subsampling that selects a specified number of ion images randomly; if a dataset has a lower number of ion images, all ion images would be considered. For each selected gold standard dataset from METASPACE and each tagger involved, we generated unique URLs containing the dataset name, the tagger name, and the maximal number of ion images to be considered. The tagging results are stored in real time, associated with the tagger name, and can be opened by the same tagger either to continue an interrupted tagging or for curation, as well as can be opened by a curator for curation; see Figure 1, video in Supplementary Data S1.

**Figure 1.**
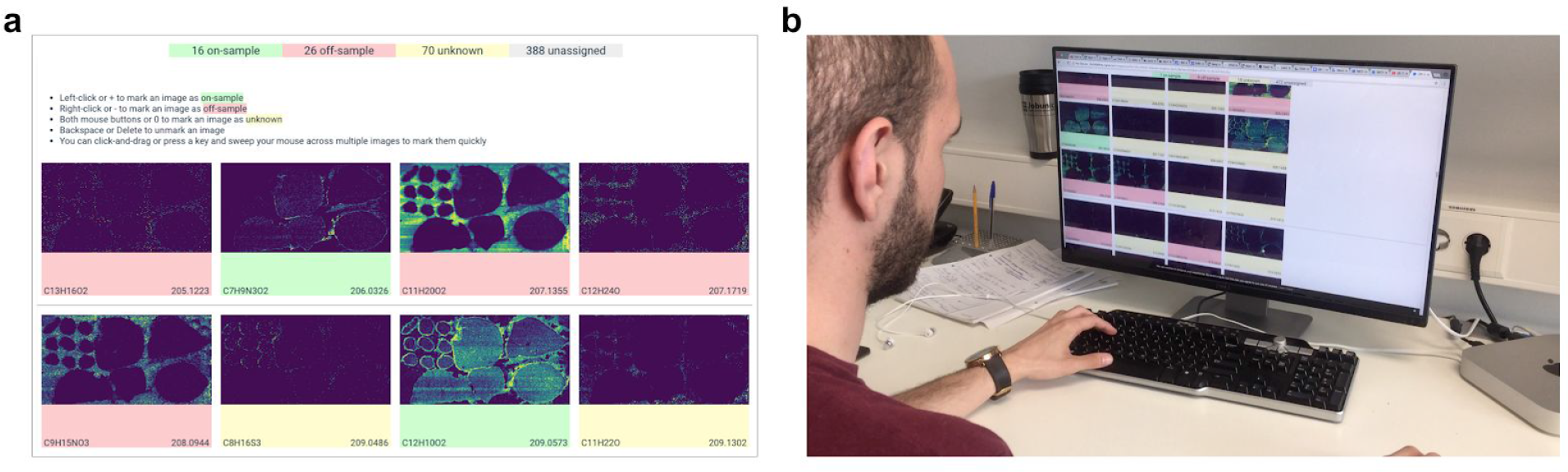
The TagOff web app for facilitated tagging off-sample ion images by using mouse clicks or keyboard shortcuts. A: The layout of the web page in TagOff. B: A tagger tagging ion images from a DESI-imaging dataset of five liver sections, contributed by Nicole Strittmatter, AstraZeneca (<u>link</u>).

#### Creating the gold standard set

For every gold standard imaging MS dataset, we considered annotations at a False Positive Rate <= 50% (the most permissive value in METASPACE) and randomly selected 500 ion images for tagging. For datasets with less than 500 ions annotated, all ion images were considered. We recruited five taggers with previous experience in mass spectrometry and explained to them the motivation behind the study. We also explained molecular and sample preparation factors contributing to generation of off-sample ion images (matrix application in MALDI, spraying in DESI), the ion suppression effect leading to lower intensities of matrix signals within the sample region, as well as advanced imaging MS effects which can affect how images look like (high noise, metabolite leakage, wet matrix application). Then, we explained to the taggers how to use the TagOff web app to tag the ion images. Importantly, the tagging was performed before algorithms for recognition of off-sample ion images were developed. This ensured that even taggers involved in the development were not biased by knowing how the final version of algorithms work.

All 87 gold standard datasets were randomly distributed between five taggers, with each dataset assigned to one tagger. Each tagger received a set of pre-generated URLs to be opened in the TagOff web app. No optical images or information about sample, imaging MS, protocols used to acquire the images were provided to the taggers. The taggers were instructed to spend less than one second per ion image.

After the tagging process was completed, “off-sample” and “on-sample” tags in each dataset were reviewed by a curator with experience in imaging MS. For each dataset, the curation consisted of a quick skimming through images of the same class and correction of the tags when necessary. The curation was intended to ensure that the tagger understood the task correctly and to minimally correct obvious mistakes. For two taggers, after seeing a high number of inconsistencies in the first 5 tagged datasets, the curator discussed it with the taggers, instructed them again and asked to repeat the tagging.

### Evaluating the gold standard

We assessed the complexity of the off-sample classification task and the reproducibility of the taggers judgements by calculating an inter-tagger agreement. For this, after the tagging was completed, the curator selected five gold standard datasets which were, in their opinion, among the most difficult datasets to tag. All taggers, who did not tag these datasets initially, were asked to tag them additionally, so that each of these five datasets was tagged by all five taggers. Then, we computed the taggers’ pairwise Cohen’s kappa agreement coefficient (Cohen, 1968) for all ion images from these five datasets which were tagged as “off-sample” or “on-sample”. The images tagged as “unknown” by at least one tagger in the pair were ignored.

### Methods for automated recognition of off-sample images

We developed three methods for automated recognition of off-sample ion images. Supplementary Data S3 includes a description of one more, template-image method, that we decided not to include in the main text.

#### Spatio-molecular biclustering method

The “spatio-molecular biclustering” method assumes that pixels of the off-sample area have similar molecular profiles as well as molecules corresponding to the off-sample area have similar distributions. First, for each dataset we found the border pixels. Note that, as discussed earlier, some of the considered datasets have non-rectangular calculated borders. Second, for each dataset we considered a matrix of ion intensities in all pixels. We performed biclustering using *SpectralCoclustering* from the scikit-learn v0.19.1. We iteratively increased the number of clusters from two to 20 until we found two large spatial clusters occupying more than *cluster_percent* (method parameter) percent of all pixels. Third, we assigned the resulting clusters to either “off-sample” or “on-sample” class. For the two large clusters, the cluster with a larger number of border pixels was assigned to “off-sample”; the other large cluster was assigned to “on-sample”. For a smaller cluster, the cluster was classified as off-sample if its border pixels occupied more than *border_percent* of all border pixels or more than *full_percent* of all small cluster pixels; otherwise it was assigned to “on-sample”. Biclustering of pixels and ions immediately provided assignment of ions to either “off-sample” or “on-sample”. Note that biclustering is an unsupervised method and requires no training. For selecting the parameters *cluster_percent*, *border_percent*, and *full_percent*, we used the five-fold cross validation. The resulting parameters were averaged over the five folds.

#### Molecular co-localization method

The “molecular co-localization” method assumes that off-sample ions are similar across datasets in terms of which ions they are co-localized with. We considered all ions annotated in at least one gold standard dataset with an FDR <= 50%. Within each dataset, we considered each ion as a vector of its intensities in all pixels. We represented all ions in a molecular space of all molecular formulas in all gold standard datasets as follows. First, we computed pairwise cosine similarities between all ions. Then, for each ion I and each molecular formula M (corresponding to a dimension of the molecular space), we considered the maximal similarity between the ion I and the ions M_n corresponding to the molecular formula M. Then, once the ions were mapped into this molecular space, we considered the same classifiers as for the “template-images” method earlier and evaluated them on the gold standard images as described later in section “Classifiers evaluation”.

#### Deep residual learning method

In the last method, we exploited Deep Residual Learning, a recently introduced deep learning approach to image recognition (He *et al.*, 2016). The advantage of the residual learning is faster training of a network which at the same time is deeper than comparable networks. Residual learning was demonstrated to outperform other deep learning approaches in the tasks of image recognition and segmentation in several competitions in 2015. We used the fastai library (v1.0.34) with the PyTorch framework (v1.0) with the ResNet-50 model from the torchvision library (v0.2.1). First, all ion images of non-rectangular datasets were padded with zeros. Then, the images were normalized with the ImageNet statistics (per channel mean and std), resized and cropped to 224×224 and, per requirement of the model, converted into RGB three-channels images by pasting the same ion image into all three channels. In order to increase the training set, we applied data augmentation with the settings default for the fast.ai library that included the following transformations: image random crop/pad, horizontal and vertical flip, image warping, rotation, zooming, brightness and contrast adjustment. We used the ResNet-50 network pre-trained on the ImageNet images (Russakovsky *et al.*, 2015). For training, we used the binary cross entropy as the loss function with the batch size of 96 and weight decay (L2 normalization) of 0.1. The training included two stages: first the training the network head (classifier) then fine-tuning the network body (feature extractors/convolutional layers). In the first stage, max learning rate (lr) was 3e-3, with five epochs. In the second stage, for the first third of the network lr was 1e-5, for the second third of the network lr was 1e-4, for the third third of the network lr was 3e-4, with five epochs. The training was done using the one cycle learning policy (see https://arxiv.org/pdf/1803.09820, by Leslie N. Smith, US Naval Research Laboratory, Washington, DC, USA) where learning rate rapidly increases to its max value and then decreases to a small value during all epochs. For the network weight optimizer we used the Adam algorithm with the max learning rate described above. The hyperparameters were selected manually by splitting the gold standard into a validation and training sets, and finding the hyperparameters providing the best F1 score for the off-sample images recognition. The validation was performed as described later.

### Classifiers evaluation

The considered classifiers were optimized to maximize the F1 score for the off-sample class. We also calculated the precision and recall to provide more detailed information about the performance. For each algorithm, we performed five-fold cross-validation when the whole gold standard was randomly split into five parts, each subsequently taken out for evaluation of the classifier trained on the rest of the data. To avoid overfitting, we used a grouped five-fold cross validation where ion images from the same dataset were all assigned either to the training or validation part.

## Results

### Selected datasets

For the gold standard set, we selected 87 public imaging MS datasets from METASPACE; see Supplementary Data S2 for a list and metadata. The datasets span various technologies and samples (see Figure 2). Importantly for the methods development, the gold standards includes datasets with both rectangular and non-rectangular imaged area. Also, some datasets included several tissue sections in the imaged area. For examples of representative off-sample ion images from the gold standard datasets, see Figure 3.

**Figure 2.**
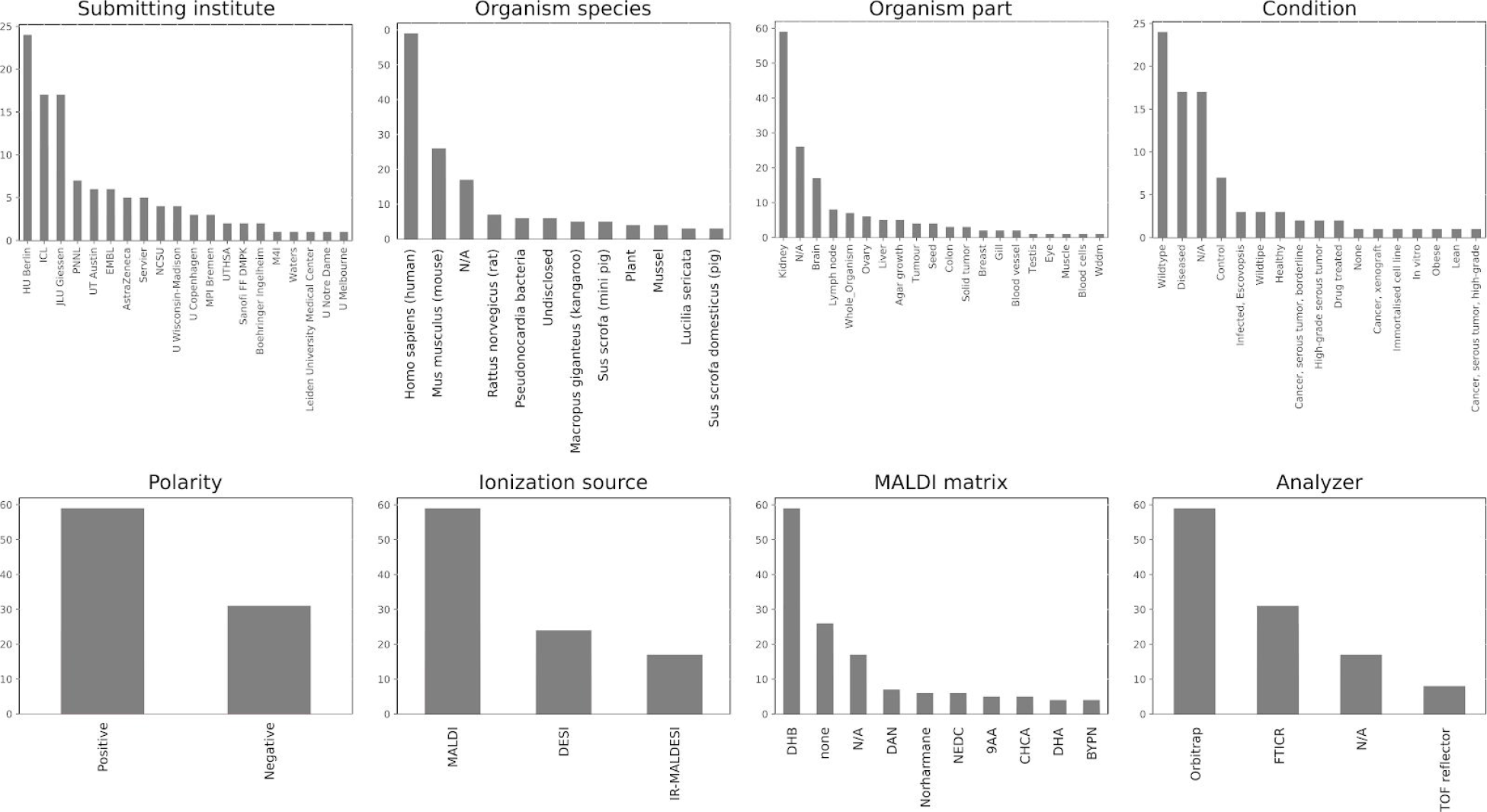
Properties of the public METASPACE datasets selected for the gold standard. They represent a variety of labs, types of samples, and technologies. Following a stratified random selection, this gold standard set reflects the full set of public datasets in the METASPACE knowledge base and, taking into account the size of METASPACE currently encompassing over 3000 public datasets, can be considered to be a representative sample in the field of imaging MS.

**Figure 3.**
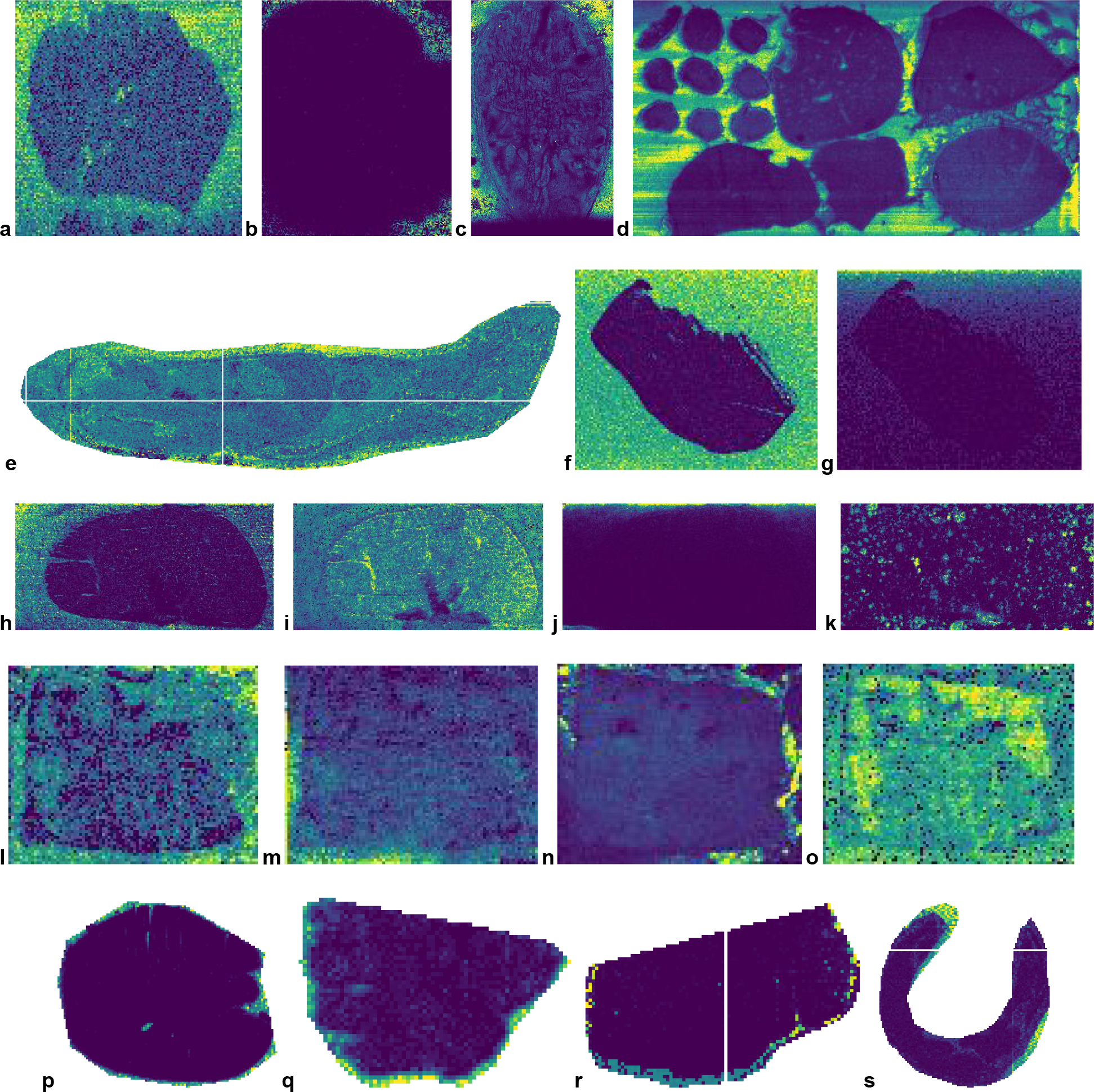
Representative off-sample ion images from the gold standard datasets illustrating a variety of spatial patterns exhibited by such images as well as particular aspects of the imaging MS datasets. A-C: Illustrative images from several datasets showing the indicative off-sample pattern (high intensities outside of the tissue section) from DESI- (A) and MALDI-imaging (B-C). D: A dataset with several tissue sections. E: A non-rectangular dataset where the acquisition area was selected around a whole body mouse tissue section. F: The indicative off-sample pattern. G: A gradually-changing off-sample pattern. H-K: Different spatial patterns of off-sample images in the same dataset (MALDI-imaging, DHB matrix). H: The indicative pattern. I: Everywhere-pattern. J: Gradually-changing pattern. K: Spotty pattern with spots everywhere, potentially due to a special matrix (DHB) application. L-O: Different spatial patterns of off-sample images in the same dataset (DESI-imaging). L: Mainly off-sample localization. M: Gradual off-sample localization. N: Off-sample leakage pattern. O: Everywhere pattern. P-S: Examples of off-sample MALDI ion images with only a narrow band of off-sample pixels due to the acquisition area selected precisely around the tissue section. METASPACE links to the ion images: <u>A</u>, <u>B</u>, <u>C</u>, <u>D</u>, <u>E</u>, <u>F</u>, <u>G</u>, <u>H</u>, <u>I</u>, <u>J</u>, <u>K</u>, <u>L</u>, <u>M</u>, <u>N</u>, <u>O</u>, <u>P</u>, <u>Q</u>, <u>R</u>, <u>S</u>. For acknowledgements for these and other gold standard datasets, see section Acknowledgements.

### Pilot study on creating a gold standard

The pilot study included tagging of thousands of ion images by two taggers (PilotTagger1 and PilotTagger2) each following their own semi-automated strategies. This experience helped us learn about a variety of spatial patterns in off-sample and on-sample images (Figure 3), complexity and subjectivity of the tagging task, pitfalls of using automated strategies, as well as estimate resources necessary to obtain a sufficiently large gold standard set of tagged ion images.

Although most off-sample ion images exhibited the indicative off-sample distribution (high intensities in the off-sample area, low intensities in the on-sample area, e.g. Figure 3a-d,f), for some datasets their off-sample ion images displayed a surprising heterogeneity in spatial patterning. Some off-sample ion images showed a gradual change of intensities (e.g. Figure 3g,j) that can be explained by spatial charging or gradual changes in calibration or intensities observed for individual molecules. Some ion images exhibited a “metabolite leakage” pattern (low intensities in the off-sample area but with high intensities at the perimeter of the section, Figure 3n) which were particularly hard and ambiguous to assign to either off- or on-sample class. Some off-sample ion images exhibited unexplainable patterns e.g Figure 3k. Some datasets contained no clear off-sample region or only a very small region containing only a narrow band of pixels around the sample e.g. Figure 3p-s. This was observed mainly for the non-rectangular datasets where the acquisition area was selected precisely around the tissue section area.

We have encountered the following issue when tagging off-sample ion images using a semi-automated strategy. We considered two semi-automated strategies. The first strategy included ranking ion images according to a particular measure and then selecting a set of images having measure values above a cutoff. The second strategy included finding a set of off-sample ion images automatically and then curating them by using a visual examination. Any of such strategies using a tagging algorithm introduce a potential bias into the gold standard. In case a classification algorithm resembles the tagging algorithm, this would lead to a misleadingly high accuracy when evaluated on such a gold standard set.

This consideration motivated us to turn to using fully-manual tagging for creating the gold standard. This required implementing a web app to facilitate tagging, as from the pilot study we learned that simple approaches not requiring any special software (e.g. manual arrangements of images into folders in a file browser or tagging of images in a spreadsheet) are not feasible for obtaining a sufficient number of tagger ion images for a gold standard for machine learning algorithms. From the pilot study, we have also learned that it is necessary to introduce the “unknown” tag because many ion images are hard to classify and assigning them to either “off” or “on” categories would be rather arbitrary and would reduce the quality of the gold standard.

### Gold standard

The obtained gold standard includes 23238 manually tagged ion images, of them 13326 “off-sample” and 9913 “on-sample”, available at https://github.com/metaspace2020/offsample. The ion images tagged as “unknown” were not included into the gold standard.

### Agreement between taggers

Table 1 shows the average pairwise inter-tagger agreement values for each of the five selected datasets. Note that those datasets, in curator opinion, were among the most difficult datasets to tag. The tagger-curator agreement values range from 0.18 to 0.69 and inter-tagger average agreement values range from −0.01 to 0.43. We speculate that the dataset “Mousebrain_MG08_2017_GruppeA” was hard to tag due to a strong sample analytes leakage into the off-sample area as well as due off-sample ion images exhibiting several spatial patterns, one of them including only a small area in the corner of the acquisition area (Supplementary Figure S1). The dataset “Servier_Ctrl_mouse_wb_median_plane_DHB” was likely hard to tag due the whole-body section including organs of various properties (density, ionization) as well as presence of the skin-localized ions which resemble matrix ions with high intensity at the perimeter of the section (Supplementary Figure S2).

**Table 1.** Agreement between five taggers and the curator for the five selected datasets from the gold standard. The datasets were selected to be among the most difficult datasets to tag. The tagger-curator average shows how well the taggers agree with the curated tags. The inter-tagger pairwise average shows the average pairwise agreement between taggers without considering the curated tags.

| **Dataset** | **Tagger-curator**<br>**average** | **Inter-tagger**<br>**pairwise average** |
| --- | --- | --- |
| 70%meoh_8cyc_75um | 0.69 | 0.39 |
| DESI quan_Swales | 0.55 | 0.43 |
| ICL//A51 CT S3-centroid | 0.41 | 0.2 |
| Mousebrain_MG08_2017_GruppeA | 0.23 | 0.12 |
| Servier_Ctrl_mouse_wb_median_plane_DHB | 0.18 | -0.01 |
| **Average** | **.41** | **.22** |

Next, we evaluated the performance of each tagger individually, and their agreement with other taggers. Supplementary Table S1 shows that taggers T1 and T2 highly agree with the curator (with the Cohen kappa >0.6), T3 and T4 moderately agree (kappa >0.3), whereas T5 has a low agreement (kappa 0.14). Note that the tagger-curator agreement closely follows the inter-tagger agreement. The opposite would indicate excessive curation that could be associated with a bias introduced by the curator.

### Performance of the off-sample recognition methods

#### Spatio-molecular biclustering method

The results of the unsupervised spatio-molecular biclustering method are shown in Table-BICLUSTERING. The optimal values of the method parameters for cluster assignment were *cluster_percent* = 0.09, *border_percent* = 0.1, *full_percent* = 0.1.

We investigated the datasets for which the method performed poorly, i.e. the F1 score per dataset for either the “off-sample” or “on-sample” class was below 0.7, which accounted to eight datasets in total. We visually inspected the spatial clusters for each dataset. For three out of eight datasets, the spatial clusters were visibly noisy (e.g. Supplementary Figure S1a) as compared to visibly homogeneous clusters for the remaining 84 datasets (e.g. Supplementary Figure S1b). For two out of poorly performed eight datasets, we observed the wrong assignment of large spatial clusters to either “off-sample” or “on-sample” class (Supplementary Figure S1c-d). We additionally examined all other datasets and confirmed that for all but these two datasets, the assignment was correct.

In order to improve the method, we considered a semi-automated strategy when a user would curate the assignment of two largest spatial clusters. This semi-automated method is easy to implement as the curation can be performed just once for a dataset, and requires a quick visual examination and one click for assignment of one of the two spatial clusters to the “off-sample” class. For the semi-automated method, we manually swapped labels for two datasets for which automated assignment failed (Supplementary Figure S1c-d) that improved the performance up to the F1 score of 0.96 (Supplementary Table S2, second row).

For 78 out of 87 datasets, the method produced two large spatial clusters per dataset. For the remaining nine datasets, two large clusters and, in addition, from one to eight smaller spatial clusters per dataset were found. In total, 14 out of 16 of the small clusters were assigned correctly.

#### Molecular co-localization method

Similar to the template-based method, the molecular co-localization method performed the best when combined with the SVM with the linear kernel, among other machine learning methods. The performance is shown in Supplementary Table S3 (first row).

Since the molecular co-localization method maps all ion images into the molecular space of all molecular formulas annotated at FDR <=50%, it is natural to assume that this method works the best when applied to datasets of similar molecular content. To test this hypothesis, we applied it only to a group of MALDI-imaging datasets acquired using the 2,5-dihydroxybenzoic acid (DHB) matrix in the positive ion mode. The polarity mode and the MALDI matrix are among the most influential parameters determining the molecular content, as molecules have preferential ionization polarity and affinity to a MALDI matrix. Supplementary Table S3 (second row) shows that indeed, the performance of the classifier has been significantly improved, particularly for recognizing off-sample images with the F1-score increased from 0.9 to 0.96. This is likely because off-sample ion images mainly correspond to matrix ions and are the most matrix-specific part of an imaging MS dataset.

#### Comparing performance of all methods

Table 2 compares performance of all developed methods for recognizing off-sample ion images. The best performing method was the deep residual learning that achieved the F1 off-sample score of 0.97. The second best method with the F1 score of 0.96 was the semi-automated spatio-molecular biclustering where curation of the cluster assignments to off/on classes was performed for two datasets. The spatio-molecular biclustering method without any curation achieved the F1 score of 0.93. Interestingly, the biclustering method is unsupervised and the gold standard was used only to select the parameters. The F1 score of the molecular co-localization method was 0.90 only. Note that the molecular co-localization method performed considerably better (F1 score of 0.96) on a reduced set of DHB positive data (Supplementary Table S3) that potentially indicates that molecular co-localization can perform on a par with other methods when applied to data collected with the same matrix.

**Table 2.**
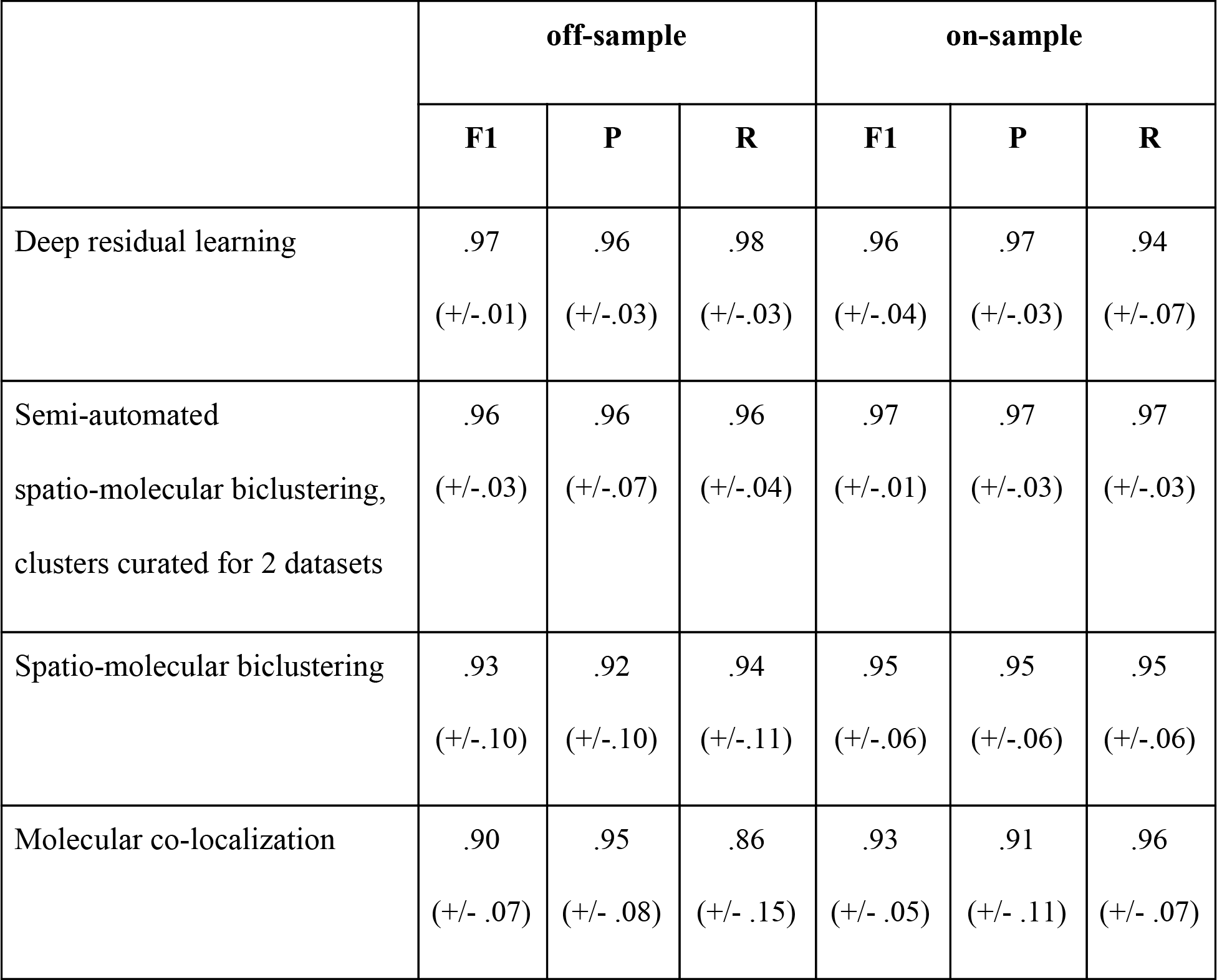
Performance of the developed methods for recognizing off-sample ion images as evaluated on the gold standard of 23238 ion images showing F1-score (F1), precision (P), and recall (R). For each measure, we show the average and confidence intervals (+- two standard deviations) over five folds of the cross validation.

Supplementary Data S3 describes the performance of the template-image method that we decided not to include in the main text which achieved the F1 score of 0.96 and 0.92 in semi- and fully-automated scenarios, respectively.

### Investigating DHB matrix clusters

Recognition of off-sample ion images can have numerous applications and generally is expected to improve downstream data analysis. Here, we decided to apply it to characterize the properties of DHB matrix clusters. For this, we selected 31 MALDI-imaging datasets from the gold standard which were acquired using the DHB matrix in the positive ion mode; see Supplementary Data S3. We generated 353 molecular formulas of the DHB matrix clusters by using a combinatorial model proposed earlier (Keller and Li, 2000). We created a custom molecular database for METASPACE by combining these molecular formulas with the HMDB v4 database (Wishart *et al.*, 2018). We added this molecular database to METASPACE and re-annotated the 31 datasets against this database.

For the 31 selected datasets, 6677 ions of DHB matrix clusters were annotated by METASPACE with an FDR <=50%. Of them, 3134 (47%) were recognized as off-sample and thus can be confidently associated with the matrix. These ions represent approximately 5% of all 58203 annotated ions for the selected datasets. Per dataset, we recognized from 0 (for one dataset only) to 314 off-sample DHB matrix cluster ions (101.1 ions on average). This quantifies the amount of biologically-unrelated information that should be removed prior to any downstream analysis.

From the recognized matrix cluster ions, 4.3% of them are isomeric with non-matrix molecules in HMDB v4. This quantifies the ambiguity of metabolite annotation which, however, turns out to be relatively low.

Figure 4 shows the parameters of the recognized matrix clusters. Interestingly, the recognized clusters exhibit broad patterns for all individual components of the combinatorial model “n*M+p*(M-H2O)-x*H+y*K+z*Na” (Keller and Li, 2000) with a slight prevalence to form clusters including two molecules of DHB. Supplementary Table S4 shows the most frequently recognized matrix clusters; see Supplementary Data S4. Two most frequent clusters were recognized in 28 out of 31 datasets. Note that the protonated ion of the intact DHB matrix, [M+H]+, the ion that one would commonly expect to be detected, was recognized only in 11 datasets (35% of all datasets). Interestingly, among the most frequently detected matrix clusters (Supplementary Table S4) there are no clusters containing potassium (K). This is also visible in Figure 4e which shows that for a prevalent majority of recognized DHB cluster ions the parameter “y” corresponding to the inclusion of potassium in the cluster ion is equal to 0. The most frequent matrix cluster with potassium was “2*M+1*(M-H2O)-1*H+1*K+0*Na”, detected in 14 datasets (45% of all datasets).

**Figure 4.**
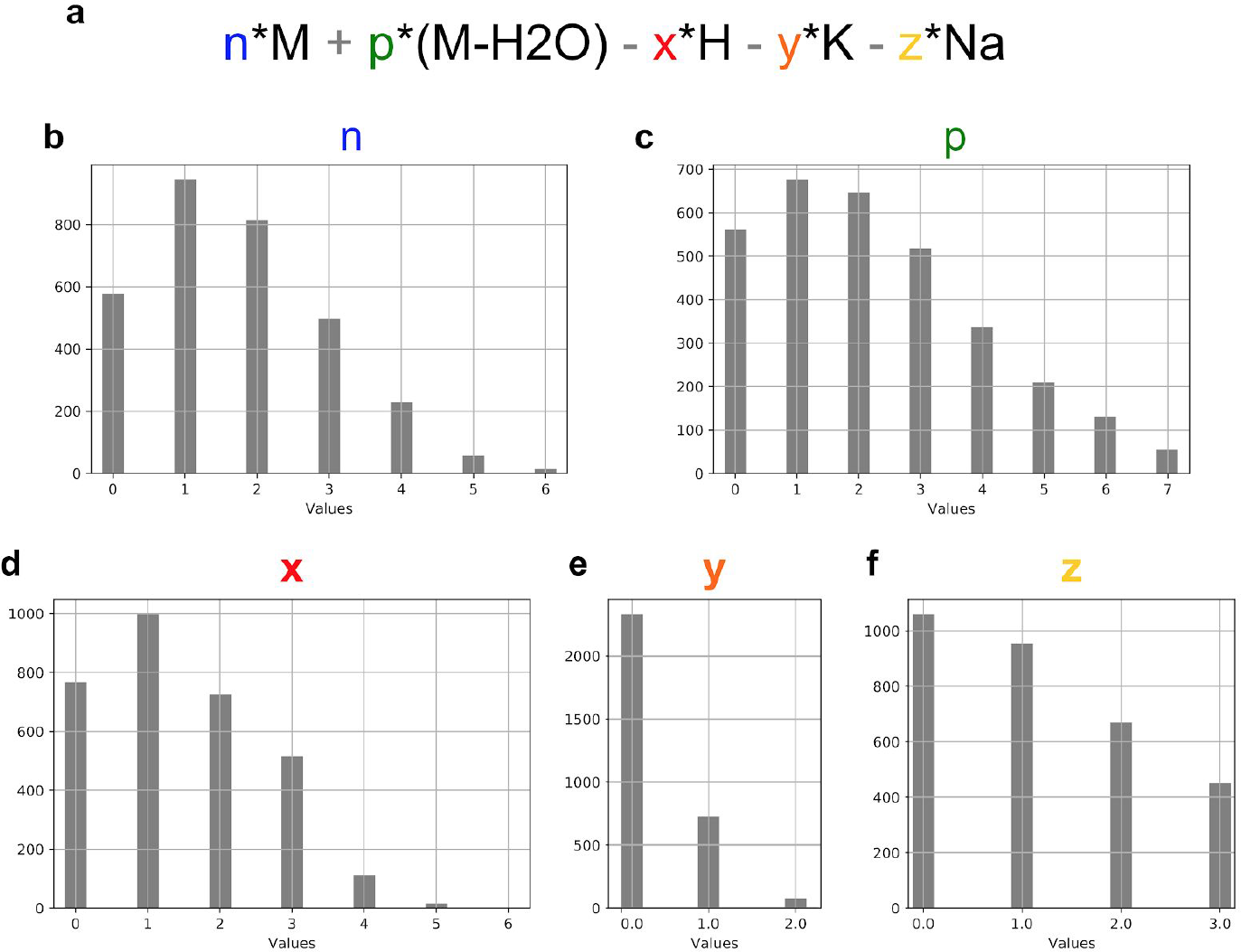
Parameters of the recognized DHB clusters. A: The formula n*M+p*(M-H2O)-x*H+y*K+z*Na represents the combinatorial model for matrix clusters from (Keller and Li, 2000) with M representing the DHB matrix molecular formula (C_7_H_6_O_4_). B-C: Histograms of parameters of the formula from A among ions in 31 MALDI-imaging DHB positive mode datasets which were annotated by METASPACE with an FDR<=50% and ion image recognized as off-sample.

## Discussion

Machine learning, and in particular deep learning, has demonstrated its capacities to outperform other approaches in various fields of science and technology including image and speech recognition, computer vision, and artificial intelligence (LeCun *et al.*, 2015). Currently, deep learning is making its way into computational biology (Angermueller *et al.*, 2016). However, for exploiting potential of machine learning algorithms, one needs annotated or tagged data for training and evaluation. When solving molecular questions in the context of imaging mass spectrometry, the annotated data is often hard to obtain due to the lack of ground truth information about molecular content of the analyzed sample. Here, we decided to explore the potential of machine and deep learning approaches to tackle a problem which on a small scale can be solved by an expert: recognition of off-sample ion images. Our expertise in creating a high-quality gold standard (Palmer *et al.*, 2015) together with the exploited techniques of modern web development helped create a high-quality gold standard of 23238 images.

It is important to highlight that this work was enabled by METASPACE, an open knowledge base of imaging MS data (http://metaspace2020.eu) (Palmer *et al.*, 2017). We used public datasets from 20 laboratories with the goal to obtain a gold standard that would be representative for the current state of the art, and to create methods that would have a wide impact in applications involving common imaging MS technologies and types of samples. By making their data public on METASPACE, these labs and their members made this study possible. Public semi-structured data is becoming increasingly useful and, through enabling novel computational developments, will ultimately benefit the field of imaging MS.

The software technologies used in METASPACE are another cornerstone in this study. The tagging web app TagOff was created based on METASPACE. Moreover, the flexible technology behind METASPACE enabled us to create a custom molecular database including DHB matrix clusters to screen for them in tens of datasets in a short time. We also used METASPACE for interactive data visualization, and sharing data and images in this paper. As a future step, we are planning to integrate the developed recognition of off-sample ion images into METASPACE and make it available to all users and in particular to those who provided public data used for training classifiers.

Working with semi-structured data from various sources is challenging because of the lack of complete information about biological systems, sample preparation, data acquisition, and data pre-processing. METASPACE metadata captures essential parts of this information (https://github.com/metaspace2020/metadata). Here, we demonstrated that the problem of recognizing off-sample ion images can be successfully solved using minimal metadata.

In this work, we compared classification approaches of different kinds: machine learning employing either molecular or spatio-molecular content as well as a computer vision approach based on deep learning. We discovered that deep learning outperforms other approaches in the F1 off-sample score, with a semi-automatic expert-driven method based on spatio-molecular biclustering almost matching and outperforming it in the F1 on-sample score. The usual drawback, also experienced by us when using various deep learning frameworks, is that deep learning networks require a substantial training time, namely days on specially-devoted GPU-equipped powerstation than hinders optimization. However, a combination of using a modern and fast framework (fast.ai), advanced deep learning methodology (residual learning), and training optimization make it possible to outperform the best expert-driven approaches.

We believe that this work will not only help solve an important problem in imaging MS, but also will serve as an example of creating a high-quality annotated gold standard and subsequently applying machine learning and deep learning methods in this field. The access to public semi-structured data allowed us to create a gold standard data which is representative for the field of imaging MS and to develop methods that will have a wide impact beyond just one laboratory or research group.

## Supplementary materials

- Supplementary Figures S1-S3
- Supplementary Tables S1-S4
- Supplementary Data S1-S5:

- **S1:** “Interactive tagging of ion images using web app.mov”, video showing a tagger using the TagOff web app to tag ion images into categories
- **S2:** “Gold standard datasets.csv”, a list and metadata of 87 public datasets from METASPACE selected for the gold standard
- **S3:** “Template-image method.pdf”, a description and results for another method not included in the main text
- **S4:** “DHB datasets.csv”, a list of 31 MALDI-imaging gold standard datasets acquired using the DHB matrix and positive ion mode selected for characterizing the properties of DHB matrix clusters
- **S5:** “DHB matrix clusters frequencies.csv”, results of annotation and off-sample recognition for 238 DHB matrix clusters generated according to a combinatorial model

## Supporting information

DHB matrix clusters frequencies

DHB datasets

Gold standard datasets

Template-image method description

Interactive tagging of ion images using web app

## Acknowledgements

We thank the image taggers: Angelos Rigopoulos, Aslihan Anal, Mohammed Shahraz, Masha Naumenko (all EMBL). We thank the contributors of all public data to METASPACE and particularly those whose data was selected for this publication: Sarah Aboulmagd, Michael Becker, Dhaka Bhandari, Mark Bokhart, Berin Boughton, Shane Ellis, Mathieu Gaudin, Erin Gemperline, Cristina Gonzalez Lopez, Richard Goodwin, Anne Mette Handler, Bram Heijs, Sophie Jacobsen, Christian Janfelt, Emrys Jones, Patrik Kadesch, Pegah Khamehgir-Silz, Mario Kompauer, Lingjun Li, Manuel Liebeke, Michael Linscheid, James McKenzie, David Muddiman, Andrew Palmer, József Pánczél, Marina Reuter, Livia S. Eberlin, Veronika Saharuka, Marta Sans, Julian Schneemann, Kumar Sharma, Bernhard Spengler, Nicole Strittmatter, Zoltan Takats, Dusan Velickovic, Eric Weaver, Guanshi Zhang. We acknowledge the funding from the EU Horizon2020 project METASPACE (No. 634402), NIH NIDDK project KPMP, ERC Consolidator project METACELL (No. 773089).

