## Supplementary Figures for "Recognizing off-sample mass spectrometry images with machine and deep learning"

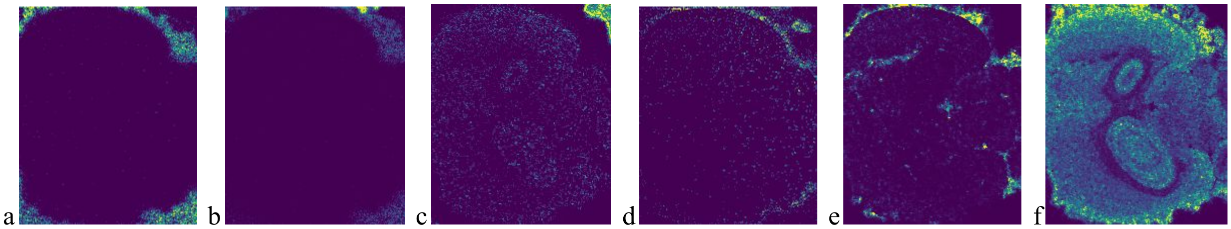


**Supplementary Figure S1.** Representative off-sample patterns for the MALDI-imaging dataset "Mousebrain_MG08_2017_GruppeA" with DHB matrix of a mouse brain section, contributed by Mario Kompauer, JLU Giessen (link). A-C show definite off-sample localizations but with different patterns. D-F represent possible sample analyte leakage into the off-sample area that is hard to assign to either off- or on-sample class without additional information. This illustrates the particular complexity of the tagging task for this dataset that was reflected in a low inter-tagger agreement (Table 1). METASPACE links to the ion images: A, B, C, D, E, F.

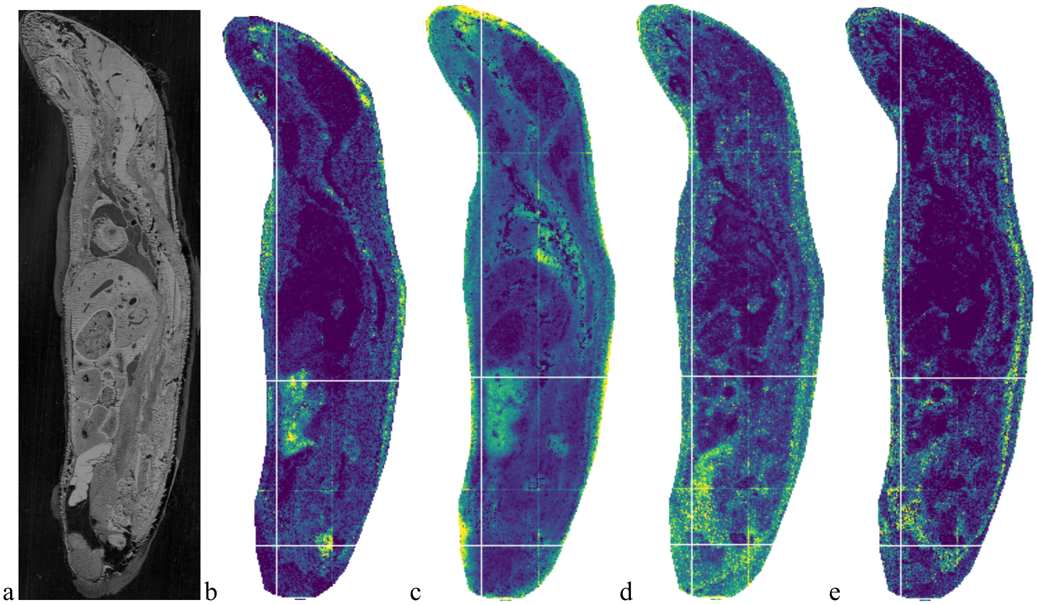


**Supplementary Figure S2.** Representative off-sample patterns for the MALDI-imaging dataset "Servier_Ctrl_mouse_wb_median_plane_DHB" from a whole-body mouse section, with DHB matrix, contributed by Mathieu Gaudin, Servier (link). **A**: Optical image of the section. B-E: Representative ion images where taggers did not agree on whether they are off- or on-sample. All ion images show some high-intensity pixels in the off-sample region but also high intensities within the tissue section, possibly due to a high complexity of the tissue section including organs of various densities and local ionization effects. Moreover, an additional complication was localization of some ions within the skin region of the whole-body section which is at the perimeter but not exactly at the boundary pixels, see in particular in B and E. This illustrates the particular complexity of the tagging task for this dataset that was reflected in a low inter-tagger agreement (Table 1). METASPACE links to the selected ion images: B, C, D, E.

a b c d

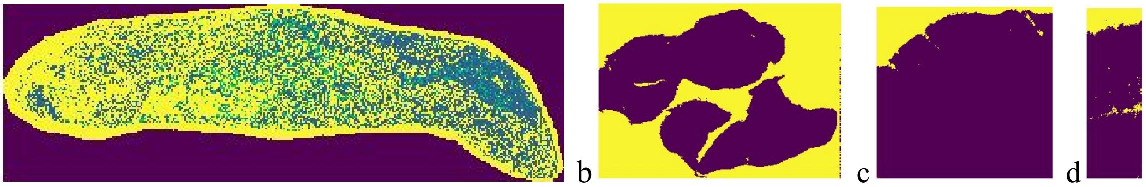

**Supplementary Figure S3. Error analysis for the spatio-molecular biclustering method. A**. An example of a noisy "on-sample" spatial cluster, MALDI-dataset of a whole-body mouse section contributed by Mathieu Gaudin, Servier (link). **B**. An example of homogeneous spatial clusters observed for a prevailing majority of datasets, DESI-imaging dataset of mouse colon with colorectal cancer contributed by Nicole Strittmatter (link). **C-D**. The datasets for which the automatic assignment of spatial clusters failed. **C.** MALDI-imaging dataset of a rat liver section, contributed by Marina Reuter, Sanofi (link). **D.** MALDI-imaging dataset of a mussel gills section, contributed by Manuel Liebeke, MPI Bremen (link).
