## Supplementary Tables for "Recognizing off-sample mass spectrometry images with machine and deep learning"

| | Tagger1 | Tagger2 | Tagger3 | Tagger4 | Tagger5 |
|---|---|---|---|---|---|
| **Tagger-curator average** | .65 | .61 | .35 | .32 | .14 |
| **Inter-tagger average** | .37 | .27 | .29 | .16 | .08 |

**Supplementary Table S1.** Detailed investigation of the five taggers with respect to their agreement with the curator as well as between each other. For each tagger, Cohen-kappa agreement of the tagger with the curator and average pairwise inter-tagger agreement is shown.

| | off-sample | | | on-sample | | |
|---|---|---|---|---|---|---|
| | F1 | P | R | F1 | P | R |
| Spatio-molecular biclustering | .93 (+/- 0.10) | .92 (+/- 0.10) | .94 (+/- 0.11) | .95 (+/- 0.06) | .95 (+/- 0.06) | .95 (+/- 0.06) |
| Semi-automated spatio-molecular biclustering, clusters curated for 2 datasets | .96 (+/- 0.03) | .96 (+/- 0.07) | .96 (+/- 0.04) | .97 (+/- 0.01) | .97 (+/- 0.03) | .97 (+/- 0.03) |

**Supplementary Table S2. Performance of the unsupervised spatio-molecular biclustering method.** F1-measure (F), precision (P), and recall (R) were calculated on the gold standard of 23238 images. For each measure, we show the average and confidence intervals (+- two standard deviations) over five folds of the cross validation.

| | off-sample | | | on-sample | | |
|---|---|---|---|---|---|---|
| | F1 | P | R | F1 | P | R |
| Molecular co-localization method, full gold standard | .90 (+/- .07) | .95 (+/- .08) | .86 (+/- .15) | .93 (+/- .05) | .91 (+/- .11) | .96 (+/- .07) |
| Molecular co-localization method, DHB positive data | .96 (+/- .06) | .96 (+/- .08) | .96 (+/- .06) | .95 (+/- .09) | .95 (+/- .07) | .94 (+/- .12) |

**Supplementary Table S3. Performance of the molecular co-localization method**. F1-score (F1), precision (P), and recall (R) are shown. The method was evaluated on the full gold standard of 23238 images, as well as on a reduced set of the gold standard MALDI-imaging datasets acquired using the 2,5-dihydroxybenzoic acid (DHB) matrix in the positive ion mode. For each measure, we show the average and confidence intervals (+- two standard deviations) over five folds of the cross validation.

| **Matrix cluster** | **Molecular formula** | **Absolute frequency** | **Relative frequency** |
|---|---|---|---|
| 1*M+2*(M-H2O)-0*H+0*K+0*Na | C21H14O10 | 28 | 90% |
| 1*M+2*(M-H2O)-1*H+0*K+1*Na | C21H13NaO10 | 28 | 90% |
| 1*M+1*(M-H2O)-0*H+0*K+0*Na | C14H10O7 | 27 | 87% |
| 1*M+1*(M-H2O)-1*H+0*K+1*Na | C14H9NaO7 | 27 | 87% |
| 1*M+1*(M-H2O)-2*H+0*K+2*Na | C14H8Na2O7 | 27 | 87% |
| 0*M+2*(M-H2O)-1*H+0*K+1*Na | C14H7NaO6 | 26 | 84% |
| 0*M+4*(M-H2O)-0*H+0*K+0*Na | C28H16O12 | 26 | 84% |
| 0*M+3*(M-H2O)-0*H+0*K+0*Na | C21H12O9 | 25 | 81% |
| 1*M+3*(M-H2O)-1*H+0*K+1*Na | C28H17NaO13 | 25 | 81% |
| 1*M+3*(M-H2O)-0*H+0*K+0*Na | C28H18O13 | 24 | 77% |

**Supplementary Table S4. Most frequently annotated and recognized DHB matrix clusters.** In the matrix cluster formula, M stands for the molecular formula of the DHB matrix ($C_7H_6O_4$). The absolute/relative frequencies stand for the number/percentage of datasets (out of 31 selected gold standard datasets) in which a particular matrix cluster was annotated by METASPACE with an FDR <=50% with an ion image recognized as off-sample.
