## Supplementary material for "Recognizing off-sample mass spectrometry images with machine and deep learning": Template-image method description

### Supplementary Data S3. Template-image method

#### Method description

The “template-based” method is based on the assumption that off-sample images within a dataset are co-localized and exhibit a relatively homogeneous distribution in the off-sample area. For each dataset, we automatically selected two template off-sample ions and two template on-sample ions. For this, we considered the border pixels of the dataset. Note that some of the considered datasets were acquired within an arbitrary (non-rectangular) area, likely to reduce the acquisition time, and thus have non-rectangular calculated borders. For each ion, we calculated the sum of its normalized intensities in the border pixels. Two ions with the maximum border intensity were selected as template off-sample ions. Two ions with the minimum border intensity were selected as template on-sample ions. We represented each ion image in the dataset as a vector of its intensities in all pixels. Then, we computed cosine similarities between each ion and the template ions. As a result, each ion was represented in the template space as a vector of the length four. Then, in this template space, we considered a variety of classifiers from the scikit-learn Python package v0.19.1, including the Nearest Neighbors, linear SVM, RBF SVM, Decision Tree, Random Forest, Naive Bayes, AdaBoost, QDA, Gaussian Process, Neural Net classifiers. We evaluated the classifiers on the gold standard images as described later in section “Classifiers evaluation”.

#### Method performance

For the template-image method, among considered classifiers applied to the low-dimensional representation of images in the space of cosine scores to the template ion images, the Support Vector Machine (SVM) classifier with the linear kernel showed the best performance. Table below (first row) shows the performance with the F1-scores equal to 0.92 and 0.95 for the off-sample and on-sample recognition, respectively.

We investigated cases when the method performed poorly. As expected, the main issue was selection of wrong templates. After visual examination of all template ions for all 87 datasets, we have identified 20 datasets for which at least one of the four automatically selected templates was incorrect as compared to gold standard annotation.

In order to improve the method, we considered a semi-automated strategy when a user would curate the selected template images. This semi-automated method is easy to implement as the operation would need to be performed just once for a dataset. For the semi-automated method, we manually selected template ions for the 20 datasets for which automated selection failed. As expected, the semi-automated method performed considerably better (see Table below, second row). We also investigated a simpler semi-automated version when only two template ions (one off-sample and one on-sample template) would be selected by a user as this version would be user friendlier in a real-life applications. The semi-automated version with two templates (see Table below, last row) outperformed the automated version and showed only a slight drop of performance as compared to the semi-automated version with four templates.

| | off-sample | | | on-sample | | |
|---|---|---|---|---|---|---|
| | **F1** | **P** | **R** | **F1** | **P** | **R** |
| Template-based method, 4 templates | .92 (+/- 0.14) | .93 (+/- 0.09) | .91 (+/- 0.20) | .95 (+/- 0.06) | .94 (+/- 0.10) | .96 (+/- 0.03) |
| Semi-automated template-based method, 4 templates | .96 (+/-.06) | .98 (+/-.02) | .94 (+/-.13) | .97 (+/-.04) | .96 (+/-.09) | .98 (+/-.01) |
| Semi-automated template-based method, 2 templates | .95 (+/-.07) | .97 (+/-.03) | .93 (+/-.14) | .96 (+/-.05) | .95 (+/-.09) | .98 (+/-.02) |

**Table. Performance of the template-based classifiers.** F1-score (F), precision (P), and recall (R) were calculated on the gold standard of 23238 images. For each measure, we show the average and confidence intervals (+- two standard deviations) over five folds of the cross validation.
